# Expanding the Landscape of Aging via Orbitrap Astral Mass Spectrometry and Tandem Mass Tag (TMT) Integration

**DOI:** 10.1101/2024.12.13.628374

**Authors:** Gregory R. Keele, Yue Dou, Seth P. Kodikara, Erin D. Jeffery, Dina Bai, Joao A. Paulo, Steven P. Gygi, Xiao Tian, Tian Zhang

**Author notes:** These authors contributed equally (Gregory R. Keele, Yue Dou).

## Abstract

Aging results in a progressive decline in physiological function due to the deterioration of essential biological processes, such as transcription and RNA splicing, ultimately increasing mortality risk. Although proteomics is emerging as a powerful tool for elucidating the molecular mechanisms of aging, existing studies are constrained by limited proteome coverage and only observe a narrow range of lifespan. To overcome these limitations, we integrated the Orbitrap Astral Mass Spectrometer with the multiplex tandem mass tag (TMT) technology to profile the proteomes of three brain tissues (cortex, hippocampus, striatum) and kidney in the C57BL/6JN mouse model, achieving quantification of 8,954 to 9,376 proteins per tissue (cumulatively 12,749 across all tissues). Our sample population represents balanced sampling across both sexes and three age groups (3, 12, and 20 months), comprising young adulthood to early late life (approximately 20-60 years of age for human lifespan). To enhance quantitative accuracy, we developed a peptide filtering strategy based on resolution and signal-to-noise thresholds. Our analysis uncovered distinct tissue-specific patterns of protein abundance, with age and sex differences in the kidney, while brain tissues exhibit notable age changes and limited sex differences. In addition, we identified both proteomic changes that are linear with age (i.e., continuous) and that have a non-linear pattern (i.e., non-continuous), revealing complex protein dynamics over the adult lifespan. Integrating our findings with early developmental proteomic data from brain tissues highlighted further divergent age-related trajectories, particularly in synaptic proteins. This study not only provides a robust data analysis workflow for TMT datasets generated using the Orbitrap Astral mass spectrometer but also expands the proteomic landscape of aging, capturing proteins with age and sex effects with unprecedented depth.

## INTRODUCTION

Progressive deterioration in fundamental biological processes results in aging-related decline, leading to increased risk of disease and mortality^1^. Loss of physiological integrity is the primary risk factor for major human pathologies, including cancer and neurodegenerative diseases^1–4^.Transcriptional correlates with aging-related decline have been well-studied, through both bulk^5^ and single-cell^6^ RNA-seq experiments. Though there have been some studies of aging in murine models^7–11^, age-related changes at the protein level are less understood, and poor correspondence with transcript aging changes has been observed in recent studies in murine tissues^12,13^. A recent discovery-based proteomic study in 10 tissues from C57BL/6JN mice revealed age differences between adult (8 months) and late midlife ages (18 months)^14^. Surprisingly, brain tissues had relatively fewer age and sex differences than other tissues. However, this study provided a limited map of age differences with only two age groups and approximately only 5,000 proteins quantified in each tissue. We hypothesized that the lack of detected age differences in brain tissues was due in part to insufficient proteome coverage and insufficient variability in surveyed age groups.

The dynamic range of the proteome is vast, with the cellular proteome spanning seven orders of magnitude^15^. Fractionation reduces sample complexity and thereby enables greater peptide and protein coverage in proteomic analysis^16,17^. However, the identification and quantitation of many low-abundance peptides still suffer from the limited sensitivity of mass spectrometers. The Asymmetric Track Lossless (Astral) analyzer enables up to 200Hz acquisition of high-resolution accurate mass (HRAM) MS/MS spectra. The parallel acquisition using the Orbitrap and Astral analyzers permits full scans with a high dynamic range and resolution and fast and sensitive HRAM MS/MS scans, expanding proteome coverage and ensuring quantitative accuracy^18–21^. Recent advances in tandem mass tag (TMT)-based proteomic technology have enabled multiplexed sample preparation and data acquisition, allowing complex experimental designs with no loss in quantitative integrity^22,23^. The high sensitivity of the Astral analyzer also enables the separation of TMTpro reporter ions and the utilization of TMTpro 18-plex for sample multiplexing. Moreover, no previous studies have showcased the use of TMT on the Orbitrap Astral Mass Spectrometer.

Here, we first integrated the Orbitrap Astral Mass Spectrometer with TMT technology to enable a multiplexed proteomic analysis with increased proteome coverage. To improve the age dimension resolution, we profiled the proteomes of three brain tissues (*i.e.*, the cortex, hippocampus and striatum) from male and female mice aged 3, 12, and 20 months, representing approximately young adulthood to early late life (approximately 20-60 years in humans). In addition to the brain tissues, the kidney with its extensively characterized sex differences^12,24^ was included. To enhance quantitative accuracy, we developed a rigorous peptide filtering strategy based on resolution and signal-to-noise thresholds and quantified approximately 9,000 proteins per tissue. Our analysis revealed distinct tissue-specific patterns of protein abundance. The dataset highlighted both age and sex differences in the kidney, while revealing age differences but limited sex differences in brain tissues. Notably, we identified both linear and non-linear (*i.e.*, continuous and non-continuous) proteomic changes with age, revealing complex protein dynamics over the lifespan. We integrated our dataset with early developmental proteomic data from brain and kidney tissues, and further highlighted divergent age-related trajectories, particularly in brain synaptic proteins. This study broadens the proteomic landscape of aging and highlights the exceptional capability of the Orbitrap Astral Mass Spectrometer to capture age-related molecular changes with remarkable depth.

## Methods

### Tissue collection

C57BL/6JN mice were ordered through NIA. Upon arrival, mice were acclimated to the local facility for one month. To collect tissues, mice were euthanized by cervical dislocation, and tissues were collected as fast as possible and flash-frozen in liquid nitrogen. Brain tissues were dissected on filter papers soaked with ice-cold PBS under a dissection microscope. Each brain region was flash-frozen in liquid nitrogen upon dissection. Kidney tissues were dissected and flash-frozen in liquid nitrogen.

### Sample preparation for proteomic analysis

Tissue samples were homogenized using the FastPrep-24 Instrument using lysing matrix D in lysis buffer (8M urea, 200 mM EPPS, Roche protease inhibitor tablets) and cells sonicated. Lysates were cleared by centrifugation (10 min at 13,000rpm at 4°C) and protein concentrations were measured using Pierce BCA assay kits. Proteins were then reduced with TCEP (5mM for 15 minutes at room temperature) and alkylated with iodoacetamide (15mM for 15 minutes in the dark). The alkylation reaction was quenched by DTT (10mM for 15 minutes). For each sample, 25µg of protein was aliquoted and diluted to a final concentration of 1mg/mL and cleaned by SP3 procedure^25^. Proteins were digested using LysC (Wako, 3 hr, 37°C, 1,400rpm on a ThermoMixer) followed by trypsin (6 hr, 37°C, 1400rpm). The resulting peptides were then labeled with TMTpro 18-plex reagents (Thermo Fisher Scientific) for 1 hour at room temperature. The reaction was quenched with hydroxylamine (final concentration of 0.5% for 15 minutes). Labeled peptides were mixed. After labeling and mixing, peptide mixtures were desalted using C18 seppak cartridges (1mg, Waters). Desalted peptides were then fractionated using basic-pH reverse phase chromatography^16^. Briefly, peptides were resuspended in Buffer A (10mM ammonium bicarbonate, 5% acetonitrile, pH 8) and separated on a linear gradient from 13% to 42% Buffer B (10mM ammonium bicarbonate, 90% acetonitrile, pH 8) over an Agilent 300Extend C18 column using an Agilent 1260 HPLC equipped with single wavelength detection at 220nm). Fractionated peptides were desalted using Stage-tips^16^ prior to LC-MS/MS analysis.

### Mass spectrometry data acquisition

Peptides were separated prior to MS/MS analysis using a Neo Vanquish (Thermo Fisher Scientific) equipped with an in-house pulled fused silica capillary column with integrated emitter packed with Accucore C18 media (Thermo). Separation was carried out with 75-minute gradients from 96% Buffer A (5% ACN, 0.125% formic acid) to 30% Buffer B (90% ACN, 0.125% formic acid). Mass spectrometric analysis was carried out on an Orbitrap Astral Mass Spectrometer (Thermo). Peptides and proteins were filtered to 1% using the rules of protein parsimony^26^. For MS1, Orbitrap resolution was set to 60k, and the normalized AGC Target was set to 200%. The purity threshold was set to 70%, and purity window is 0.5%. For MS2, the isolation window was set to 0.5 m/z. HCD collision energies were set to 35%, with a maximum injection time is 20ms. FAIMS voltages of -30, -35, -45, -55, -60 were used.

### Mass spectrometry data analysis

Spectra were converted to mzXML using a modified version of ReAdW.exe. Database searching included all entries from the *Mus musculus* UniProt database (January 26, 2022). This database was concatenated with one composed of all protein sequences in the reversed order. Searches were performed using a 50-ppm precursor ion tolerance for total protein level profiling. The product ion tolerance was set to 0.03 Da. These wide mass tolerance windows were chosen to maximize sensitivity in conjunction with COMET searches and linear discriminant analysis^27,28^. TMT tags on lysine residues and peptide N termini (+304.2071Da) and carbamidomethylation of cysteine residues (+57.021 Da) were set as static modifications, while oxidation of methionine residues (+15.995 Da) was set as a variable modification. Peptide-spectrum matches (PSMs) were adjusted to a 1% false discovery rate (FDR)^29,30^ . PSM filtering was performed using a linear discriminant analysis, as described previously^27^ and then assembled further to a final protein-level FDR of 1%^29^. TMT reporter quantitation was performed using both 0.001 Da and 0.003 Da peak match tolerance (PTM) windows. Dataset using the 0.001 Da PTM was used for the subsequent analysis. 0.001Da Peptide-Spectrum Matches (PSM) with any of the 18 reporter ions resolutions lower than 45K and a total signal-to-noise (S/N) ratio less than 1,440 across 18 channels were filtered out. Proteins were quantified by summing reporter ion counts across all matching PSMs, as described previously^31^. The signal-to-noise (S/N) measurements of peptides assigned to each protein were summed and these values were normalized so that the sum of the signal for all proteins in each channel was equivalent to account for equal protein loading.

### Peptide filtration and summarization to proteins

Peptides that did not have full resolution and/or S/N lower than 1440 for 18 samples were removed and not used for protein analysis. Abundance levels for proteins (based on UniProt ID) were estimated by summing their retained peptides. The 18-plex TMT study designs simplified data processing and normalization because all samples per tissue could be analyzed within a single batch and thus not requiring a batch adjustment.

### Detecting age- and sex-related differences for proteins in individual tissues

For a given tissue, we tested for proteins that differed across age groups, fit either as continuous or non-continuous (categorical), sexes, and age-by-sex groups. To test for age differences, we fit the following log-linear regression model for each protein:

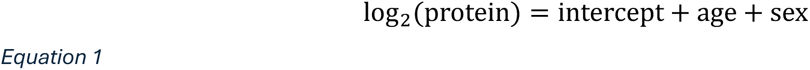

where age represents either a continuous covariate (in terms of months), and thus the model estimates only a single coefficient or a non-continuous categorical covariate factor, fit as two coefficients. The age difference was then tested with an F-test by comparing to a null model excluding the age term. Sex was tested similarly, except with the null model excluding the sex term instead. To test for age-by-sex differences, we fit the following regression model:

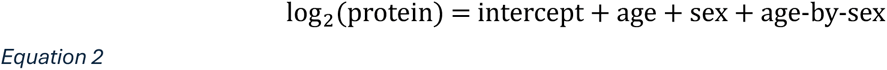

where age-by-sex represents the interaction effect between age and sex (with age fit as either a continuous or non-continuous term). The interaction term was then tested using an F-test, comparing the model in Eq. 2 to a null model excluding the interaction term (Eq. 1). The p-values were then FDR-adjusted using the Benjamini-Hochberg method^32^.

We also performed these tests for individual peptide level data to assess trends for how resolution and S/N affected coefficient estimation when compared to protein-level coefficients (**Fig. S1**). To assess whether the age differences were more consistent with a continuous fit or non-continuous fit, we compared the non-nested models using the Bayesian information criterion (BIC), a model selection criterion. A lower BIC corresponds to the BIC-supported model.

### Detecting consistent and distinct age- and sex-related differences for proteins across brain tissues

We jointly modeled tissues together to detect age differences that were consistent across tissues, representing the main effect of age, as well as allowing age-by-tissue differences to capture tissue-distinct differences. We expanded the model in Eq. 1 to:

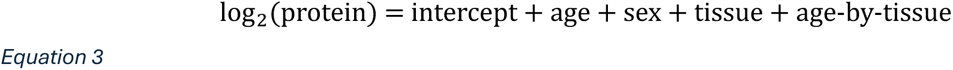

where the model now includes a main effect for tissue and its interaction with age. We tested for the age main effect with an F test comparing Eq. 3 to a null model fit excluding the age term. We also tested for an age-by-tissue effect with an F test comparing Eq. 3 to a null model fit excluding the interaction term. The same approach was also used to test for a sex main effect and sex-by-tissue interaction across tissues. The p-values were then FDR-adjusted using the Benjamini-Hochberg method^32^.

### Principal component analysis (PCA)

PCA was performed using the PCA R package^33^. We combined data across all tissues, representing 70 samples, after filtering to the 5778 proteins observed in all four tissues. We first log_2_ transformed each protein to account for an expected overall log-linear distribution. We estimated 10 principal components (PCs).

### Gene set enrichment analysis (GSEA)

GSEA was performed using the clusterProfiler R package^34^. Age- and sex-differences were used to define gene sets (*e.g.*, age coefficient > 0 and P_adj_ < 0.1 for a given tissue), which were then compared to a universe gene set (*e.g.*, all analyzed proteins for a given tissue). Enriched gene ontologies and KEGG pathways were detected based on and FDR threshold, such as P_adj_ < 0.1. We also defined gene sets based on stepwise categorical age trends (*e.g.*, “Down-Flat” for a protein that did significantly dropped between 3 and 12 months but did not change between 12 and 18 months). We were more stringent in defining age trend gene sets (FDR < 1%) given the greater potential to over-fit the data. The unique gene identifiers were ENSEMBL gene IDs for gene ontology analysis and UniProt IDs for KEGG pathway analysis.

### Analysis of kidney and brain proteomic data from Wang *et al.* 2024^11^

We re-analyzed the kidney and brain tissue data from Wang *et al.* 2024^11^ to allow for consistent processing and modeling with the present study. This study was a cross-sectional study of early development age groups of one week, one month, and two months. After filtering out proteins with > 80% missing values for a tissue, we used the same statistical modeling of age- and sex-differences as with the current study, including both continuous and non-continuous age modeling (Wang *et al.* 2024 used non-continuous modeling in their reporting^11^). We also performed PCA and GSEA as described for the current study.

### Software

All data processing and statistical analyses related to age differences were conducted with the R statistical programming language^35^.

## RESULTS

For each tissue, three replicates of male and female C57BL/6JN mice at the ages of 3, 12, and 20 months were multiplexed in a TMTpro 18-plex (**Table S1**). The samples were fractionated into 96 fractions using offline HPLC and recombined into 24 samples^16^. Twelve out of 24 samples were then analyzed using the Orbitrap Astral Mass Spectrometer (**Fig. 1A**).

**Figure 1.**
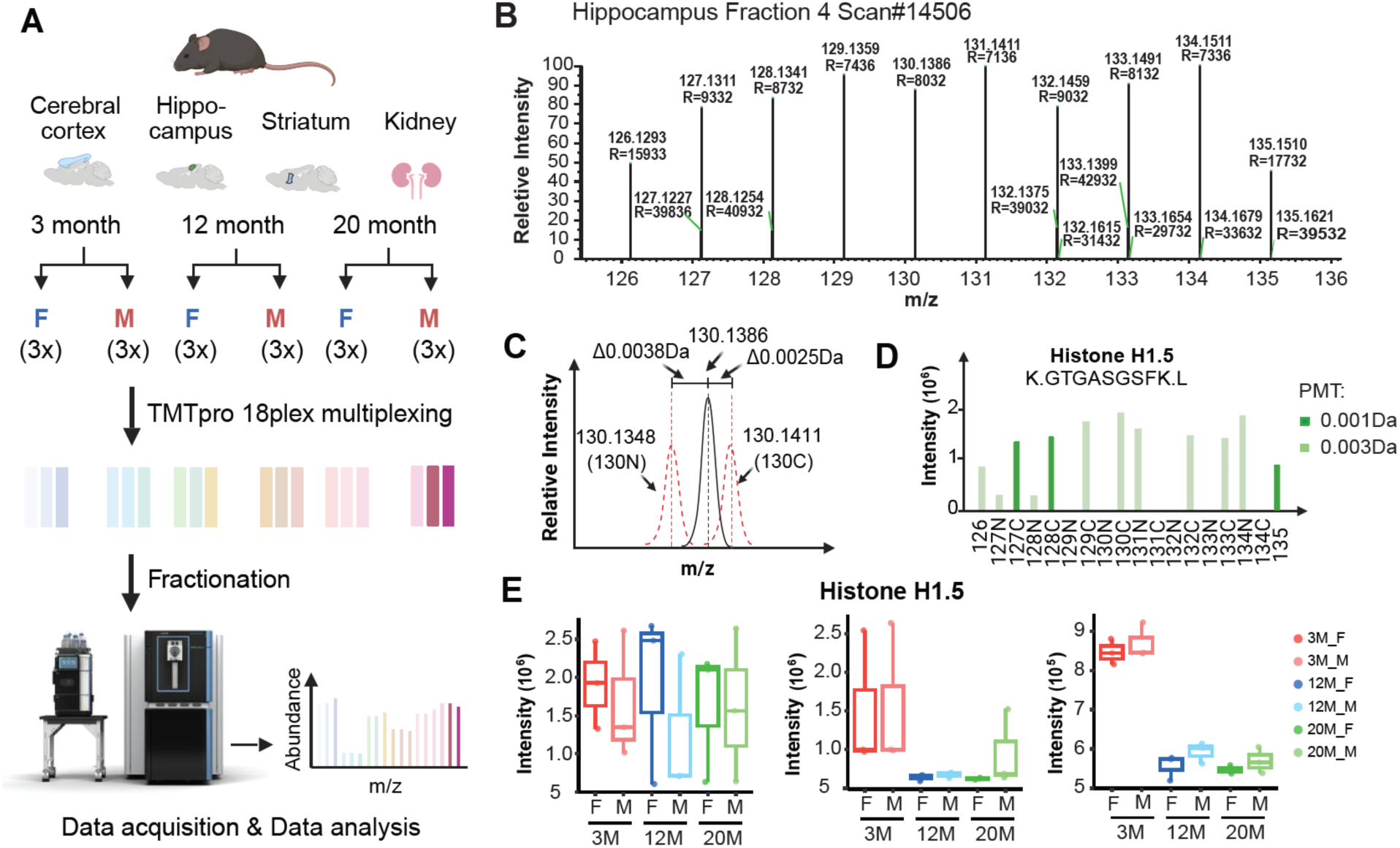
Peptide Filtering Enhances TMT Quantification Accuracy with Orbitrap Astral Mass Spectrometry. **(A)** Overview of the workflow. For each tissue, three male and female C57BL/6JN mice at ages 3, 12, and 20 months were multiplexed. Samples were digested, labelled using TMTpro 18plex, combined, fractionated and recombined into 24 fractions, of which 12 were analyzed on an Orbitrap Astral Mass Spectrometer. (B) Reporter ion region of a peptide, GTGASGSF, from Histone 1.5, in the hippocampus tissue is shown. (C) Ion saturation leads to inaccurate measurement of TMT reporter ions. The peak at m/z 130.1368 shifted by 0.0038 Da from 130N and 0.0025 Da from 131C due to ion saturation, showing coalescence within a narrow range. (D) Quantitative impact of saturated ions on peptide using different peak match tolerance (PTM) windows. A PMT window of 0.001 Da for quantitation enabled detection of 3 reporter ions (dark green) compared to 12 reporter ions (light green) detected with a 0.003 Da window, though it did not correct for the inaccurate quantitation of this peptide. (E) Excluding this peptide from the analysis corrected for the distortion in Histone 1.5 protein quantitation, revealing a significant age effect. The left panel shows protein quantitation with the peptide using PMT 0.003 Da, while PMT 0.001 Da was used for the middle panel. The right panel shows protein quantitation after removing the peptide with ion saturation effects; PMT 0.001Da was used.

### Enhancing TMT Quantification Accuracy with Orbitrap Astral Mass Spectrometry through Peptide Filtering

From these data, we noticed that the peptides with high intensities can yield erroneous results if their ion intensities reach 10^6^, leading to distorted and non-ideal detector response. Reporter ion intensities from a single scan for the peptide GTGASGSF from Histone 1.5 are shown as an example (**Fig. 1B-D**). Instead of 18 reporter ions, 12 peaks were detected in the reporter ion m/z range, with twelve of the carbon and nitrogen isotope versions of the TMT tag coalesced into six peaks (**Fig. 1B**). As an example, the peak merged identified with m/z 130.1386 is 0.0038Da to the right of 130N and 0.0025Da to the left of 131C (**Fig. 1C**). Using 0.001 Da peak match tolerance (PTM) window for reporter ion quantitation resulted in the quantitation of 3 reporter ions (dark green) compared to 12 reporter ions using 0.003 Da PTM window. However, this adjustment failed to mitigate the impact of the inaccurate peptide quantitation on the protein quantitation (**Fig. 1D**). This erroneous quantitation of this peptide led to a distortion of the quantitation of Histone 1.5. Removing this peptide from protein quantitation revealed an age effect on protein H1.5 (**Fig. 1E**). Detector saturation caused by high-intensity ions (i.e., ion saturation) has been widely reported in time-of-flight (TOF) mass spectrometers and has also been observed in other mass spectrometry platforms, including triple quadrupoles and trap-based instruments^36^. In this study, we hypothesized that the low resolution resulting from ion saturation contributed to the inaccurate quantification of TMT reporter ions.

To systematically investigate the saturation effects in these datasets, we developed a C# program to extract the resolution of each TMT reporter ion in each scan. The program recorded the resolution, intensity, and noise of each reporter ion with a peak match tolerance of 0.001 Da for each MS2 spectrum from the raw file, enabling detailed downstream analysis. Subsequently, the resolutions of the 18 TMT reporter ions associated with peptides used for protein quantitation were analyzed (**Table S2**). A resolution of 45K is required to distinguish the carbon and nitrogen isotopologs of the TMT tags, which have a mass difference of 0.006 Da^23,31^. To assess the impact of resolution, the log10 ratio of the total signal-to-noise (S/N) across 18 channels for each MS2 spectrum was plotted for the four tissue types, focusing on cases where any of the 18 reporter ions fell below the 45K resolution threshold (**Fig. 2A**). The results revealed a bimodal distribution in resolution corresponding to low and high S/N groups. Less than full resolution (*i.e.*, poor resolution) in the low-S/N group was attributed to low-intensity ions, whereas poor resolution in the high-S/N group was attributed to ion saturation. We investigated further whether detector saturation was an issue in TMT datasets collected using Orbitrap analyzer. We downloaded raw files from PXD018886^37^ and PXD032843^38^, checked the MS2 and MS3 data collected in Orbitrap analyzer, and detected no PSMs with saturated effects, indicating that this is a specific issue to the Orbitrap Astral analyzer.

**Figure 2.**
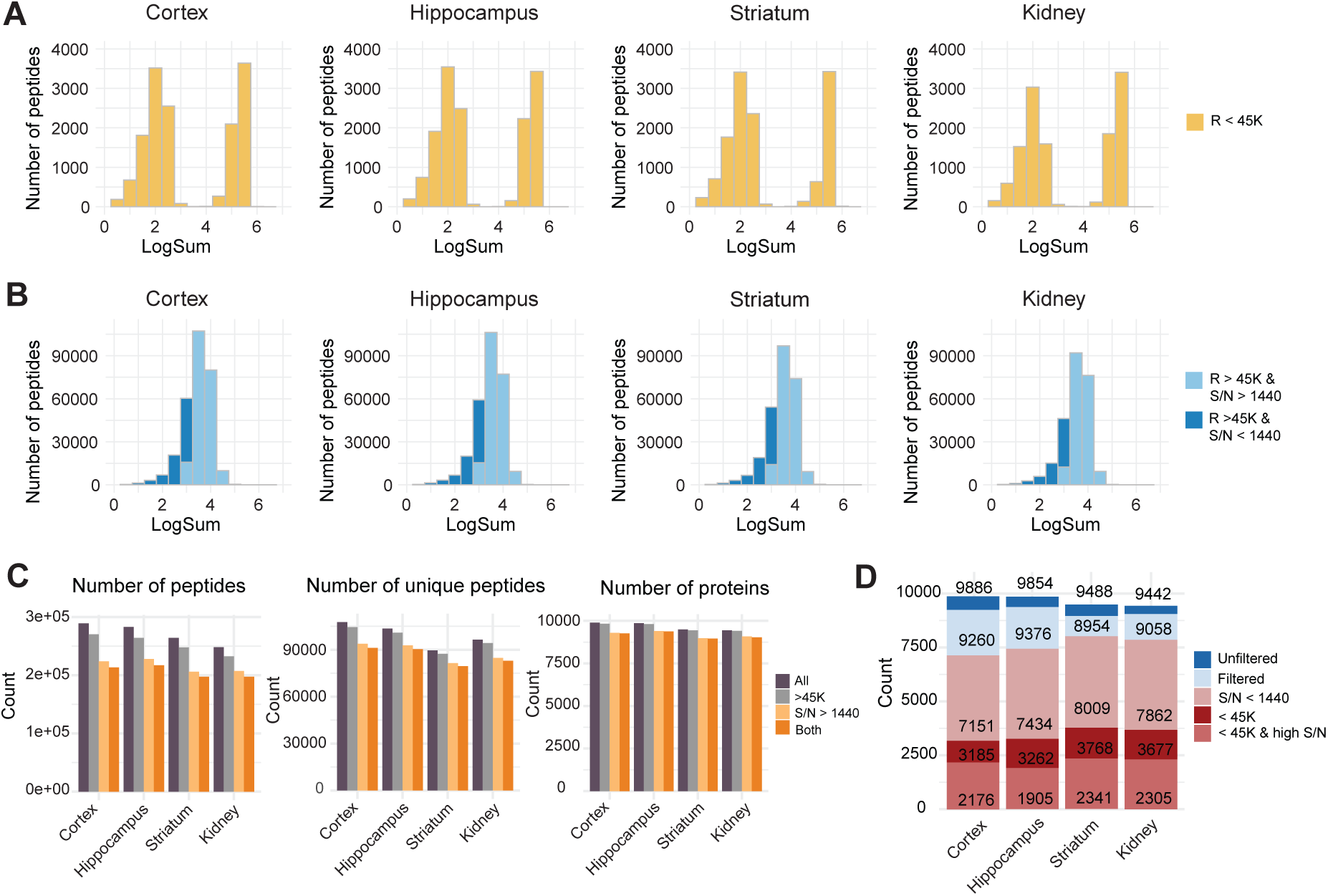
Impact of resolution and Signal-to-Noise (S/N) Filtering on peptide and protein quantitation. (A) The log_10_ ratio of the total S/N across 18 channels for each MS2 spectrum with any resolution below the 45K threshold was plotted for four tissues, showing a bi-modal distribution for low and high S/N. (B) The intensity distribution of each MS2 spectrum with low S/N after resolution-based filtering. (C) Resolution and S/N filtering effects on peptide and protein quantification. Approximately 6% of PSMs were excluded due to resolution filtering, while ∼20% were removed due to low abundance. These exclusions resulted in a 2.5% reduction in unique peptides for resolution filtering and a 9%-10% reduction for low-abundance filtering. Consequently, the number of quantified proteins decreased by approximately 5%. (D) Impact of filtering on protein quantification. Resolution filtering affected over one-third of all quantified proteins, while saturated ions impacted more than 20%. Additionally, S/N filtering affected approximately 80% of the quantified proteins.

To assess how resolution and signal-to-noise (S/N) could influence the detection of quantitative differences due to biological factors of interest, we tested for differences between age groups and sex using regression (Methods) in peptides and proteins (summed from their component peptides). Peptides with low resolutions and low intensities were associated with significant differences (*i.e.*, effects) (**Fig. S1A-B**). Comparing the regression coefficient for a protein to the coefficients for its peptides revealed a low correlation for low S/N peptides, which had coefficients closer to zero and thus would likely bias protein abundance towards the null of no detectable differences (**Fig. S1C-F**). The measurement of low-intensity peptides is more likely to be compromised by noise and interference. Excluding these peptides with low resolutions and low intensities improves the accuracy of protein quantitation and facilitates the detection of biologically relevant quantitative differences ^39^.

To avoid bias from low-S/N and low-resolution peptides, we estimated abundance levels for each protein from its component peptides after filtering out peptides that did not have full resolution and S/N lower than 1440 for 18 samples. The protein abundance estimate for an individual was then the sum of their S/N levels for the high confidence peptides. Approximately 6% of all the PSMs were removed by resolution filtering, while ∼ 20% of all the PSMs were removed due to being of low intensity. The resolution and intensity filtering led to approximately 2.5% and 9%-10% drop of the number of unique peptides, respectively (**Fig. 2B-C**). Although the filtering did not result in a dramatic drop, with an approximate 5% reduction in protein numbers (**Fig. 2C**), the impact on protein quantitation was substantial, affecting a large number of proteins. More than 1/3 of all the quantified proteins were impacted by the resolution filtering, with saturated ions impacting greater than 20% of all quantified proteins and the S/N filtering impacting ∼80% of all the quantified proteins (**Fig. 2D**).

After filtering, our dataset resulted in a higher level of coverage at both the peptide and protein levels compared to TMT datasets collected on other mass spectrometers^14,16,37^, with quantitation of 9,260 proteins in cortex, 9,376 in hippocampus, 8,954 in striatum, and 9,058 in kidney (**Table S3, Fig. S2A**). Across the four tissues, 12,749 proteins were quantified. The most common overlap/unique category in the UpSet plot was 5,778 proteins observed in all four tissues, followed by 1,588 in just kidney and 1,541 in all three brain tissues (**Fig. S2B**). These numbers are higher (by ∼1,000 or more per tissue) than recent aging protein studies in mice^10,11,14^, which included both data-independent acquisition (DIA) and other tandem mass tag (TMT) experiments.

### Tissue, age, and sex are major drivers of protein abundance variation

We next sought to characterize variation in protein abundance across tissues, age groups, and sexes. Examining correlation at the level of tissue type, we observed strong a correlation between the brain tissues (**Fig. S3A**). The brain tissues were less correlated with kidney tissue, which displayed notable sex-based variation (**Fig. S3A**).

We then performed principal component analysis (PCA) to determine whether age and sex drive the observed variation (Methods). The first principal component (PC1) primarily distinguished the brain tissues from kidney (**Fig. S3B**). PC2 primarily distinguished the striatum from the other tissues and PC3 distinguished the hippocampus and cortex from each other and the other tissues (**Fig. S3B-C**). Cumulatively, tissue type explained 97.5% of the variation (**Fig. S3D**). PC4 (0.4%) aligned cleanly with sex, which was strongest in the kidney as expected^12,14^ (**Fig. S3E**). Age correlated with PCs 5 and 6, representing 0.5% of the overall variation (**Fig. S3F**). These results support the potential for tissue-specific signals as well as the large-scale presence of sex and age differences at the level of individual proteins.

### Age differences for individual proteins are highly prevalent in brain tissues

To identify proteins with age and sex effects, we tested for individual proteins with abundance differences between age groups, sexes, and age-by-sex groups within a single tissue (Methods, **Table S4**). We first modeled age as both a continuous term, thus estimating a single coefficient and assuming a continuous (*i.e.,* linear) trend with age.

The number of proteins with continuous age differences ranged from 1,109 (in hippocampus) to 4,196 (in striatum) (**Fig. 3A, S2C**). In the brain, sex differences were relatively minor, ranging from 41 (in cortex) to 242 (in striatum) (**Fig. 3B, S2C**). Kidney had 2,242 proteins with age differences and notably 4272 with sex differences. Y chromosome-encoded EIF2S3Y and GSTA4 were shown as examples (**Fig. S2D-E**). Gene set enrichment analysis^34^ (GSEA) was performed using gene sets defined by continuous age differences (increasing and decreasing), age trends categories, and sex differences (increased in males or females) are provided in **Tables S5-S8**, which include results based on gene ontologies and KEGG pathways. GSEA findings in kidney included increased abundance with age for immune-related proteins (**Fig. S4**) and confirmation of previous signals^24^ such as increased ribosomal proteins in males and large-scale metabolic differences between the sexes. GSEA results for the brain tissues revealed a general pattern of increases in metabolic proteins and decrease in synaptic proteins. Cortex was an outlier based on having over 3,000 categorical age-by-sex differences (**Fig. S2C**), exemplified by USP24 (**Fig. S2F**). Overall, these proteins were enriched in functions related to endoplasmic reticulum, synapses, and neurons more broadly (**Table S5**).

**Figure 3.**
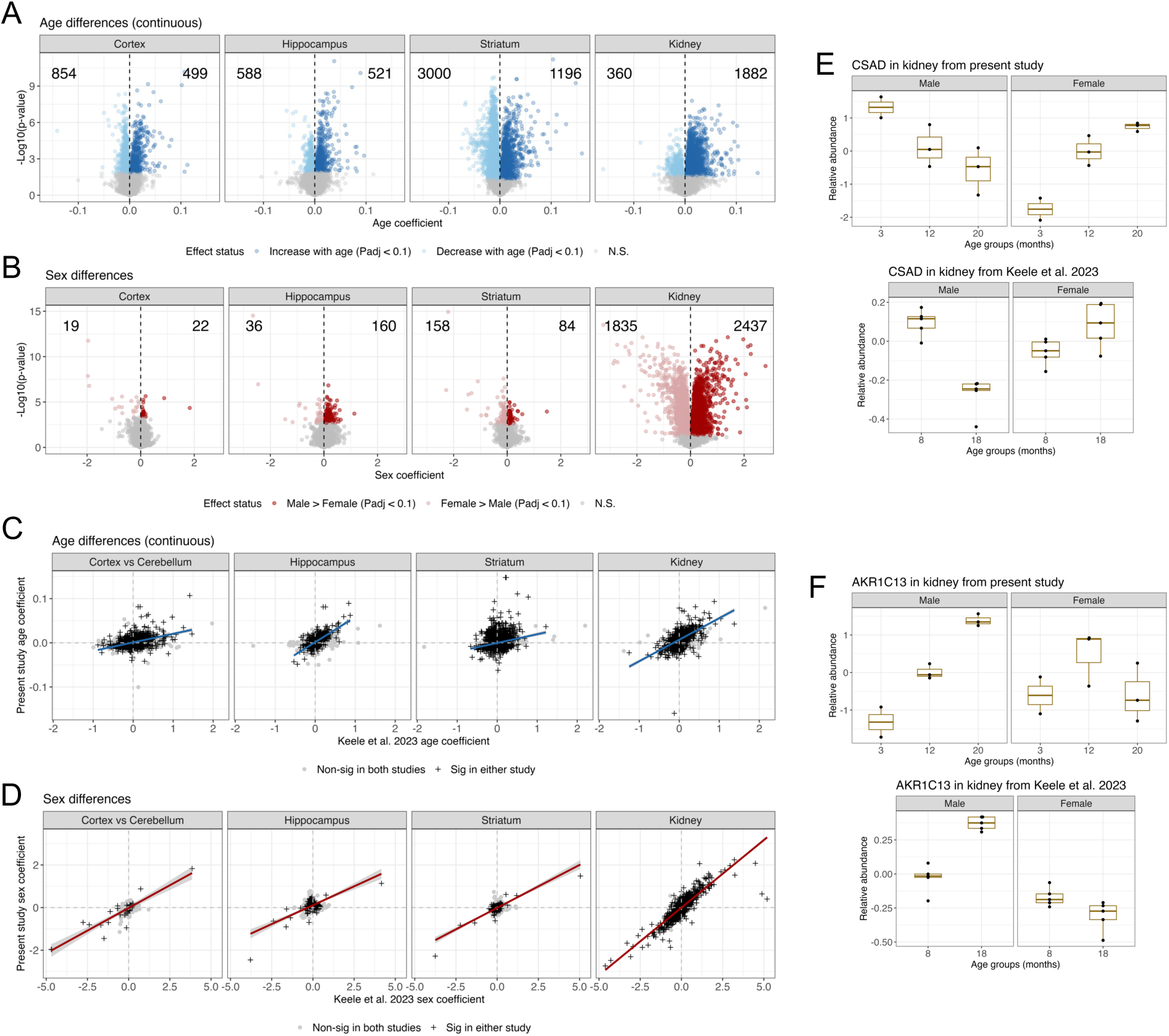
Age and sex differences detected in present study and comparison to Keele *et al.* 2023. (A) Volcano plots of continuous age differences for four tissues. Counts of statistically significant differences (FDR < 0.1) included for both directions of coefficient. (B) Volcano plots of sex differences for four tissues. Counts of statistically significant differences (FDR < 0.1) included for both directions of coefficient. (C) Comparison of continuous age difference coefficients between present study and Keele *et al.* 2023. Best fit regression line and standard error interval based on the coefficients that were significant in both studies (P_adj_ < 0.1) included for reference. (D) Comparison of sex difference coefficients between present study and Keele *et al.* 2023. Best fit regression line and standard error interval based on the coefficients that were significant in both studies (P_adj_ < 0.1) included for reference. (E) Boxplots of CSAD in kidney, which had a consistent age-by-sex difference in the present study (top) and Keele *et al.* 2023 (bottom). Data were scaled prior to plotting. (F) Boxplots of AKR1C13 in kidney, which had a consistent age-by-sex difference in the present study (top) and Keele *et al.* 2023 (bottom). Data were scaled prior to plotting.

We compared age and sex coefficients from our study to those reported by Keele *et al.* 2023^14^, which also employed a TMT study design but utilized an Orbitrap Fusion Lumos Tribrid mass spectrometer and included only two age groups: 8 and 18 months-old. Another distinction was cerebellum was profiled rather than cortex. Keele *et al.* 2023^14^ only detected 10 proteins in hippocampus, 0 protein in striatum and 496 proteins in kidney tissue with age differences. To test whether the greater number of proteins with age differences in our study were systematically driven by the increased detection of low-abundance proteins, we compared the S/N of proteins unique to our study with those quantified in both studies. We found that most proteins detected with age effects in our study were quantified in both studies. Furthermore, the proteins observed in only this study had lower average S/N and a lower rate of age differences detected compared to proteins observed in both studies (**Fig. S5**). These findings suggest that the identification of more proteins with age effects in our study is likely a result of improved statistical power due to observing a greater dynamic range of ages (three age groups rather than two). Despite the difference in the number of proteins with detected age differences, we observed consistent age coefficients for proteins with detected differences in either study (**Fig. 3C**). Similarly, sex differences were also highly consistent in the kidney, as well as for the few that were detected in the brain tissues in either study (**Fig. 3D**).

We then leverage these two studies to validate complex age-by-sex differences as demonstrated for proteins such as CSAD (**Fig. 3E**) and AKR1C13 (**Fig. 3F**) in the kidney. CSAD exhibited a consistently increasing trend in females but a decreasing trend in males across the age groups. In contrast, AKR1C13 showed a steady increase in males, while females displayed a non-monotonic up-down pattern, which only becomes evident with the broader age range. The inclusion of three distinct age groups in our study provided the resolution to capture non-linear age dynamics that would be difficult to detect with only two age groups. The observation of consistent interaction effects across independent studies strongly supports the biological validity of these findings.

### Detecting non-continuous age differences in individual proteins

Age and sex effects may not follow a proportional increase with age^40,41^, and non-linear or non-continuous patterns in these effects could be missed in our linear regression model. To investigate proteins exhibiting non-continuous effects and evaluate whether the data were more consistent with a continuous or non-continuous age difference, we expanded our approach to model categorical (*i.e.*, non-continuous) age trends using Bayesian information criterion (BIC) to compare non-nested models (Methods). Briefly, a lower BIC represents a better model fit; we categorized age differences as continuous or non-continuous based on the model with a lower BIC. Overall, we observed more continuous age differences than non-continuous (**Fig. 4A**). Each brain tissue has more than twice as many continuous age differences (continuous to non-continuous ratios ranging from 2.24 to 3.56), whereas the kidney has a lower rate of continuous age differences comparatively (continuous to non-continuous ratio of 1.24).

**Figure 4.**
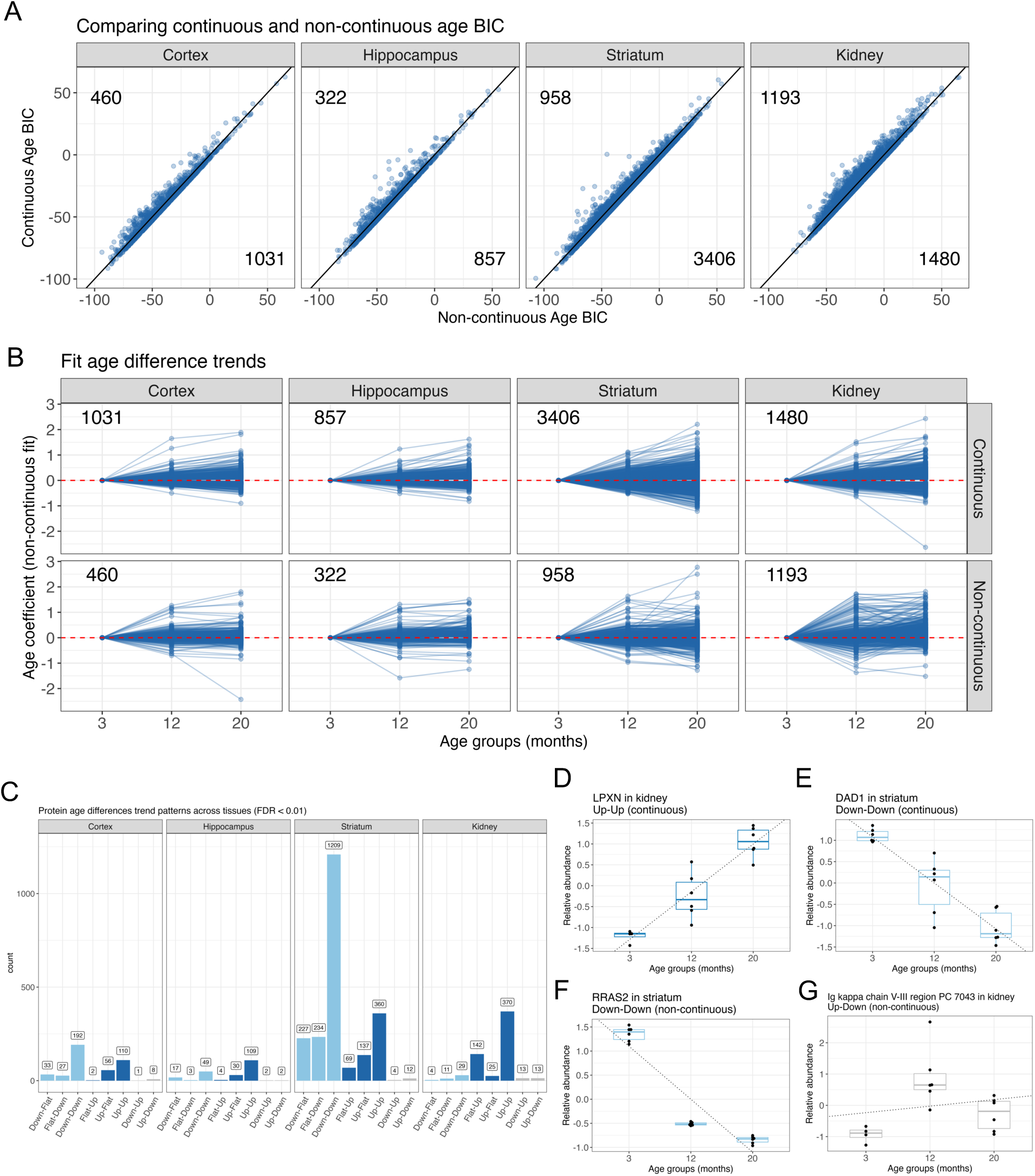
Proteins with distinct aging trends observed across tissues. (A) BIC comparison of continuous and non-continuous model fits of the age term for significant age differences (FDR < 0.1). Counts of proteins more consistent with a continuous age model are in the bottom right and a non-continuous model are in the top left. (B) Age trends for proteins with significant age differences (FDR < 0.1). Points represent age coefficients from a non-continuous model fit. Trends are categorized based on whether a continuous or non-continuous fit was preferred by BIC. (C) Categorization of highly significant (FDR < 0.01) age trends as stepwise changes between age groups. (D) Boxplots of LPXN in kidney, which had an Up-Up (continuous) trend with age. Dotted continuous bet fit line included for reference. Data were scaled prior to plotting. (E) Boxplots of DAD1 in striatum, which had a Down-Down (continuous) trend with age. Dotted continuous bet fit line included for reference. Data were scaled prior to plotting. (F) Boxplots of RRAS2 in striatum, which had a Down-Down (non-continuous) trend with age. Dotted continuous bet fit line included for reference. Data were scaled prior to plotting. (G) Boxplots of Ig kappa chain V-III PC 7043 in kidney, which had an Up-Down (non-continuous) trend with age. Dotted continuous bet fit line included for reference. Data were scaled prior to plotting.

We further categorized age trends based on stepwise changes between adjacent timepoints by testing the specific timepoint effect in the categorical age model. For example, a trend of “Up-up” represents a protein that has increased abundance at 12 months relative to 3 months and increased abundance for 20 months relative to 12 months. In all tissues, the most prevalent trend categories were “Down-Down” and “Up-Up”, which is consistent with the higher prevalence of continuous age differences. Examples of proteins with different stepwise age trends include LPXN’s increasing continuous trend in kidney, DAD1’s decreasing continuous trend in striatum, Rras2’s decreasing but non-continuous trend in striatum, and an immunoglobulin variable chain’s “Up-Down” trend in kidney (**Fig. 4C-F**). We performed GSEA for gene sets defined by categorical age trends (**Tables S5-S8**). Generally, the strongest enrichment signal was found for “Down-Down” and “Up-Up” trends, consistent with the patterns observed with continuous age (synapse-related proteins and metabolism, respectively). Some of the age trends that are less consistent with a continuous age pattern potentially reveal new aging dynamics, such as Rac protein signaling and specific immune ontologies (viral response and T cell mediated) in striatum proteins with “Flat-Up” trends (**Table S7**). These represent systematic age changes that appear to occur primarily between mid to late adult life.

### Detecting protein age difference patterns across brain tissues

To investigate tissue-specific and non-tissue-specific age and sex effects, we performed hierarchical clustering using 5,778 proteins detected in all four tissues. This analysis revealed that the brain tissues clustered together based on both sex and age differences, while the kidney tissue exhibited a distinct pattern (**Fig. 5A-B**). Building on this, we focused on data from the brain tissues to identify age-related differences that were either consistent across tissues or unique to specific ones. We filtered for the 7,377 proteins detected in all three brain tissues and tested for age-related differences, as well as age-by-tissue interactions (Methods, **Table S9**). This analysis allowed us to pinpoint proteins with either uniform age trends or tissue-specific variations (Methods).

**Figure 5.**
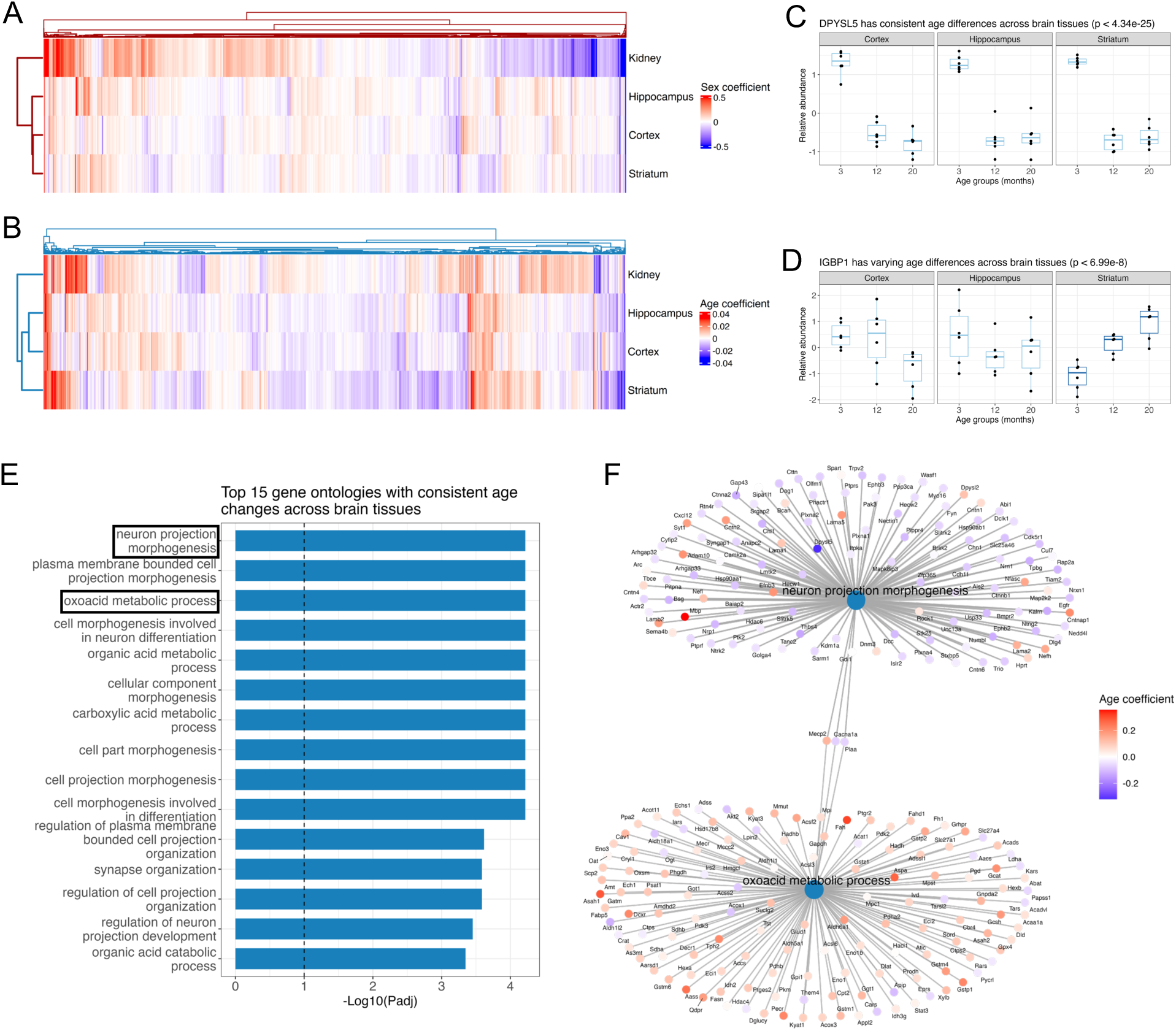
Cross-tissue modeling of age and sex differences. (A) Heatmap of sex difference coefficients for 5778 proteins observed in all four tissues. (B) Heatmap of continuous age difference coefficients for 5778 proteins observed in all four tissues. (C) Boxplots of DPYSL5, which had consistent age trends across the three brain tissues. Data were scaled per tissue prior to plotting. (D) Boxplots of IGBP1, which had unique age trends across the three brain tissues. Data were scaled per tissue prior to plotting. (E) Top 15 gene ontologies enriched in proteins with consistent age differences across the three brain tissues (FDR < 0.1). The vertical dashed line represents a threshold of P_adj_ < 0.1. Neuron projection morphology and oxoacid metabolic process are highlighted with black boxes, which are featured in Fig. 5F. (F) Network plot of GSEA results for proteins with consistent age trends across the three brain tissues.

This analysis also enabled us to identify proteins with consistent age trends across brain tissues, as well as those with tissue-specific variations. For example, we identified DPYSL5, which displayed a consistent non-continuous age trend across all brain tissues, and IGBP1, which exhibited an increasing continuous age trend specific to the striatum (**Fig. 5C-D**). We performed GSEA^34^ based on the set of 1,235 genes with evidence for consistent age differences (marginal age P_adj_ < 0.01 and age-by-tissue P_adj_ > 0.2) to identify gene ontologies associated with shared aging patterns across the brain. We observed overarching patterns of proteins related to metabolism (*e.g.*, oxoacid metabolic process) increasing with age and neurons and synapses (*e.g.*, neuron projection morphogenesis) decreasing with age (**Fig. 5E-F**). The observed negative aging trends for proteins involved in synapse biology aligns with previously reported findings by Tsumagari *et al.* 2023^10^, which profiled the cortex and hippocampus proteomes of male C57BL/6 Jcl mice in very similar age groups (3, 15, and 24 months-old). Notably, we confirmed this trend in both male and female mice in our study.

### Comparing protein age dynamics between adulthood and early development

While most age-related differences appeared continuously across the span of adulthood studied here, some proteins exhibited non-continuous trends. For instance, AKR1C13 showed a decreasing trend in females during middle-to-late adulthood (**Fig. 3F**). However, this non-continuous pattern became apparent only when young adult female mice were included in the analysis. This observation led us to explore how protein age differences during adulthood in our study compare to changes observed in early life development. To address this, we integrated our findings with early developmental data from Wang et al^11^. This complementary study (**Fig. 6A**) profiled the proteomes of 10 tissues, including whole brain and kidney, in both female and male C57BL/6JN mice across infancy-to-adolescent stages (1 week, 1 month, and 2 months). To compare between the studies, we re-analyzed the brain and kidney data from Wang et al ^11^ using a consistent statistical approach to minimize methodological differences (Methods.). Fundamental differences remain between the studies (*e.g.*, different institutions at different times, sister strains of mice, differences in tissue collection, and DIA versus TMT proteomics). PCA revealed similarities and differences in the distribution of variation across the two datasets. In Wang *et al*.^11^, the first three principal components accounted for 91.8% of the total variation (**Fig. S6A**), compared to 97.5% in the present study (**Fig. S3D**). Despite this, the contributions of tissue, age, and sex to variation differed slightly between the studies. In Wang et al., PC1 primarily separated brain and kidney, while PC2 distinctly separated age groups (**Fig. S6B**), with the youngest age group (1 week) showing a marked separation from the older age groups (1 and 2 months). Sex differences, on the other hand, strongly correlated with PC5 (**Fig. S6C**). Interestingly, compared to adulthood in the present study, age explained a larger proportion of overall variation during early development, whereas sex differences were more prominent in adulthood. Additionally, age-related changes during early development appeared less continuous (**Fig. 6B**) compared to the smoother age trends observed in adulthood (**Fig. 4A**).

**Figure 6.**
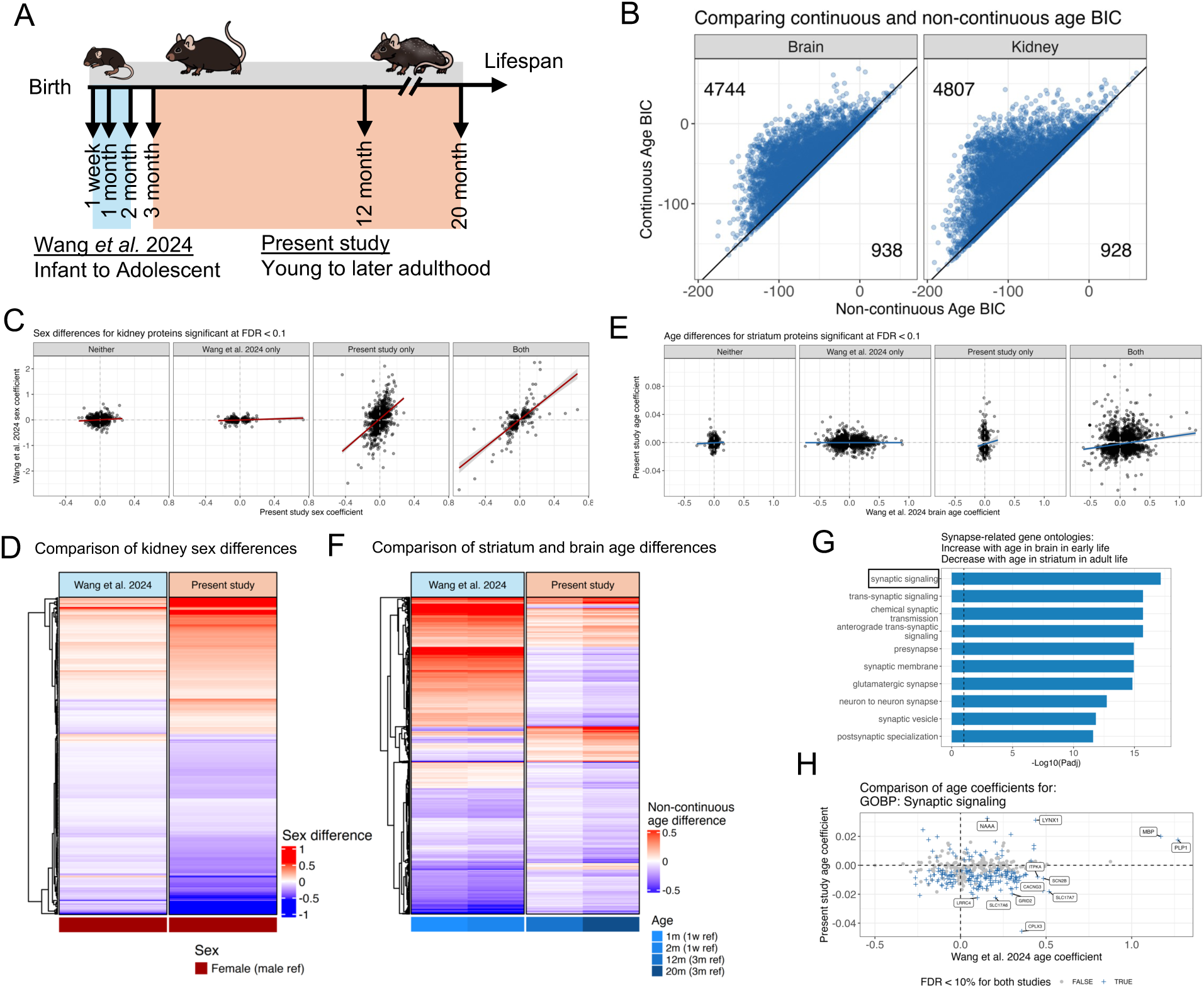
Comparison to early development protein data from Wang *et al.* 2024 reveals distinct aging patterns from adulthood. (A) Diagram of the age groups observed across Wang *et al.* 2024 and present study. (B) BIC comparison of continuous and non-continuous model fits of the age term for significant age differences (FDR < 0.1). Counts of proteins more consistent with a continuous age model are in the bottom right and a non-continuous model are in the top left. (C) Comparison of sex difference coefficients for kidney between the present study and Wang *et al.* 2024. Best fit regression line and standard error interval included for reference. Coefficients are stratified into groups based on which studies detected a significant difference (FDR < 0.1). (D) Heatmap of sex difference coefficients for kidney proteins that were significant (FDR < 0.1) in both present study and Wang et al. 2024. (E) Comparison of age difference coefficients between the present study’s striatum tissue and Wang *et al.* 2024’s brain tissue. Best fit regression line and standard error interval included for reference. Coefficients are stratified into groups based on which studies detected a significant difference (FDR < 0.1). (F) Heatmap of non-continuous age difference coefficients that were significant (FDR < 0.1) in both present study striatum proteins and Wang et al. 2024 brain proteins. Black box highlights proteins with increasing trends with age in early development and decreasing trends in adulthood, which are enriched in synapse-related functions. (G) Top 15 gene ontologies enriched in proteins with upward age trends in brain tissue during early development and downward age trends in striatum through adulthood (FDR < 0.1). The vertical dashed line represents a threshold of P_adj_ < 0.1. Vesical-mediated transport in synapse and synaptic vesical cycle are highlighted with black boxes, which are featured in Figure 6H. (H) Comparison of age difference coefficients between the present study’s striatum tissue and Wang *et al.* 2024’s brain tissue for proteins in the synaptic signaling gene ontology. Vertical and horizontal dashed lines included for reference, highlighting proteins with discordant age coefficients between the two studies

### Protein sex differences in kidney become more pronounced in adulthood

We next compared sex and age difference coefficients between the two studies. In the kidney, we observed weak concordance for age coefficients (**Fig. S7A-B**) but strong concordance for sex coefficients between early development and adulthood, particularly if the difference was statistically significant (FDR < 10%) in both studies (**Fig. 6C-D**). Interestingly, sex differences detected only in the adulthood data from the present study showed moderate concordance with early development data, while differences unique to the earlier developmental stages exhibited much lower concordance. These findings suggest that sex differences are initiated early in development but become more pronounced in adulthood, potentially coinciding with sexual maturity. In the brain, sex differences were relatively sparse in both studies. Moreover, those identified differences were largely inconsistent in the direction of the coefficients between early development and adulthood (**Fig. S7C**), suggesting a more complex or variable progression of sex differences in brain tissues across the lifespan.

### Complex age dynamics of synapse-related proteins throughout lifespan in brain tissues

We then compared the Wang *et al.* 2024^11^ brain data to all three brain tissues (cortex, hippocampus, and striatum) in the present study (striatum in **Fig. 6E**; all brain tissues in **Fig. S7D**). We note that the continuous age coefficient is a less accurate summary for the extensive non-continuous age differences of early development. Notably, far more age differences were detected in early development, and these did not strongly correlate with adulthood age differences. Even when a protein’s age difference was statistically significant in both studies (FDR < 10%), many showed discordant patterns, with the direction of change often reversed between early development and adulthood (**Fig. 6F** for striatum). One notable pattern within proteins exhibiting a potential reversal over the lifespan—specifically an increase in abundance during early development paired with a decrease in adulthood—were enriched for genes associated with synapse biology (**Fig. 6G-H**). This observation is highly consistent with synapse density across lifespan as measured through deep synaptome profiling^42^, and implies that activating the developmental network in the old brain may be an effective strategy to counteract age-related structural changes and functional decline.

## DISCUSSION

In this study, we integrated the Orbitrap Astral Mass Spectrometer with TMT technology for a multiplexed proteomic analysis. This approach enabled us to overcome previous limitations in proteome coverage and depth and boost the quantified protein number significantly, yielding insights into tissue-specific, age-related, and sex-based protein abundance patterns. During our analysis, we discovered that ion saturation in the Astral analyzer during TMT reporter ion quantitation can lead to inaccuracies, potentially resulting in the false discovery of biological effects. This issue is specific to the TMT and Orbitrap Astral combination and does not affect other DDA or DIA applications using other mass spectrometers. It can be mitigated by injecting less material; however, this approach may reduce the depth of protein quantitation. To address the challenge, we developed and applied a stringent peptide filtering strategy based on resolution and signal-to-noise (S/N) thresholds specifically tailored for the TMT datasets generated on the Orbitrap Astral system. Although the filtering process resulted in a 20% reduction in the total number of peptides and a 15% decrease in unique peptides, the impact on protein quantitation was minimal, with only a ∼5% reduction in the total number of quantified proteins (**Fig. 2C**). However, the filtering process influenced a significant proportion of proteins. Resolution-based filtering affected more than one-third of all quantified proteins, while saturated ions impacted over 20%, and S/N filtering impacted approximately 80% of the quantified proteins (**Fig. 2D**). Despite these reductions, the filtering strategy effectively mitigated biases associated with low-abundance and low-resolution peptides. This improvement in data quality enhanced the accuracy of protein quantitation, enabling us to capture complex molecular changes linked to aging with greater confidence. Our strategy provides the proteomics community with a robust approach for analyzing TMT datasets generated with the Orbitrap Astral Mass Spectrometer.

We profiled both sexes across three distinct age groups (3, 12, and 20 months) and provided a comprehensive view of proteomic changes from young adulthood to early late life. We observed distinct tissue-based proteomic trends, notably the sex differences in the kidney as contrast to the minimal sex influence in brain tissues^24^. This finding aligns with the known susceptibility of the kidney to sex-linked aging effects. Compared to previous studies, the wider dynamic range of ages included in this study improved statistical power and enabled the identification of more proteins with sex and age differences (**Fig. S5**).

From a discovery perspective, this study also identified compelling examples that can be further investigated to elucidate underlying aging mechanisms. One such example is AKR1C13, which exhibits a non-monotonic up-down pattern in female mice, a trend that only becomes apparent with our broader age range (**Fig. 3F**). AKR1C13 is an NADP(H)-dependent oxidoreductases belonging to the aldo-keto reductase (AKR) family^43^; It has broad substrate specificity for 20α-, 17β- and 3α-hydroxysteroids, and non-steroidal alcohols^44^. The increased level of AKR1C13 in adult female mice may indicate a heightened demand for lipid metabolism. In contrast, the decreased level of AKR1C13 in aged female mice could contribute to the accumulation of lipid peroxidation products, elevated reactive oxygen species (ROS), and increased oxidative damage^45^. This study functions as a quantitative protein resource for the aging-focused research community. We have made our data and processed results available to the aging research community, accessible online. It allows for querying of proteins of interest, facilitating deeper exploration of the dataset.

Prior aging studies have revealed complex non-continuous transcriptional aging trends, such as Schaum *et al.* 2020^5^, which profiled 10 age groups. Having three adult age groups in the present study and three early development age groups from Wang *et al.* 2024^11^, allowed us to more deeply characterize age trends than previous aging proteomic studies. We observed the presence of non-continuous age-related protein trajectories, emphasizing the complexity of aging at the molecular level. Expanding the dynamic range of age within a study to broadly encompass the entire murine lifespan would enable more complex models that capture highly non-continuous dynamics. Based on three age groups per sample population, the minimum number of groups to assess deviation from a continuous trend, our comparisons of developmental and adulthood aging dynamics revealed at times opposite trajectories, such as for proteins involved in synaptic functions. This highlighted that the aging process not only involves a linear accumulation of changes but also complex reversals and non-linear patterns across the lifespan. Our power to capture these patterns is limited when comparing independent studies and motivates future studies with more age groups and tissues within a single study.

Overall, this work provides a robust analysis framework for TMT datasets generated using Orbitrap Astral mass spectrometer, illustrating the potential of this advanced proteomic technology for mapping age-related molecular alterations across tissues. We expanded the proteomic landscape of aging by increasing the extent of protein quantitation across a wider range of lifespan, highlighting the need for further research to explore these age-related protein dynamics, particularly the roles of synaptic proteins in the aging processes and age-associated disease conditions.

### Limitations

While this study provides valuable insights into age-associated proteomic changes, it also has some limitations. First, our analysis was limited to four tissues. Expanding the study to a broader range of tissues could provide a more comprehensive view of proteomic aging processes across the body. Second, we applied a stringent cutoff to filter low-intensity ions, which, while necessary for ensuring quantitation accuracy, may have excluded biologically relevant peptides. This approach might result in a minor loss of depth and could overlook subtle yet meaningful proteomic variations that contribute to the aging process. Finally, this study focused primarily on protein abundance changes. Including post-translational modification (PTM) analysis in future research would offer a deeper regulatory perspective on aging, providing insights into protein functionality and interaction that abundance data alone cannot capture. Future studies addressing these limitations could build on these findings to achieve a more integrated and translatable understanding of proteomic aging dynamics.

### Data availability

The mass spectrometry proteomics data have been deposited to the ProteomeXchange Consortium via the PRIDE [1] partner repository with the dataset identifier PXD058160 and 10.6019/PXD058160. Log in to the PRIDE website using the following details: Project accession: PXD058160; Token: TMDUSpfYvdKf. Alternatively, reviewers can access the dataset by logging in to the PRIDE website using the following account details: Username;Password: fko8eHs1nW8Q. All processed data and R code to perform statistical analyses and generate the reported findings are publicly available at figshare (10.6084/m9.figshare.28016501).

## TABLES

**Table S1. Mouse sample information.** Spreadsheets linking TMT tag to mouse sample Information (age group, sex, individual identifier) for each tissue.

**Table S2. Peptide signal-to-noise information.** Spreadsheets providing quantitative signal-to-noise data for all TMT tags for each peptide in each tissue. Additional information includes peptide sequence, gene symbol, ENSEMBL gene and protein identifiers, resolution, and cumulative signal-to-noise.

**Table S3. Protein signal-to-noise information.** Spreadsheets providing quantitative abundance (signal-to-noise) data for all TMT tags for each protein in each tissue. Additional information includes gene symbol and ENSEMBL gene and protein identifiers.

**Table S4. Association summary statistics for individual tissues.** Spreadsheet providing results from statistical testing of differences in protein abundance based on sex, continuous age, non-continuous age, and the interaction between sex and age, for each tissue.

**Table S5. GSEA results from cortex.** Gene sets were defined based on statistically significant age or sex difference and the direction or type of difference pattern.

**Table S6. GSEA results from hippocampus.** Gene sets were defined based on statistically significant age or sex differences.

**Table S7. GSEA results from striatum.** Gene sets were defined based on statistically significant age or sex differences.

**Table S8. GSEA results from kidney.** Gene sets were defined based on statistically significant age or sex differences.

**Table S9. Association summary statistics for joint modeling of brain tissues.** Spreadsheet providing results from statistical testing of differences in protein abundance based on sex, continuous age, non-continuous age, and the interaction between sex and age, jointly across brain tissues.

## Acknowledgements

We thank all the Zhang Lab members for active discussion and support. T.Z. was supported by NCI grant R00CA273170. X.T. was supported by NIA grant R00AG068303. We thank Corinne M. Keele for her illustrations of a B6 mouse across its lifespan (**Fig. 6A**).

## Author contributions

X.T., and T.Z. conceptualized the project and designed the experiments. Y.D., S.P.K., and T.Z. performed the experiments. E.D.J., J.A.P., S.P.G. and T.Z. designed the methodology. G.R.K. designed the statistical approach. G.R.K., D.B. and T.Z. performed the analysis. G.R.K., J.A.P., S.P.G, X.T. and T.Z. wrote the manuscript. All authors reviewed the manuscript.

## Inclusion and diversity

We support inclusive, diverse, and equitable conduct of research.

---

**Figure S1.**
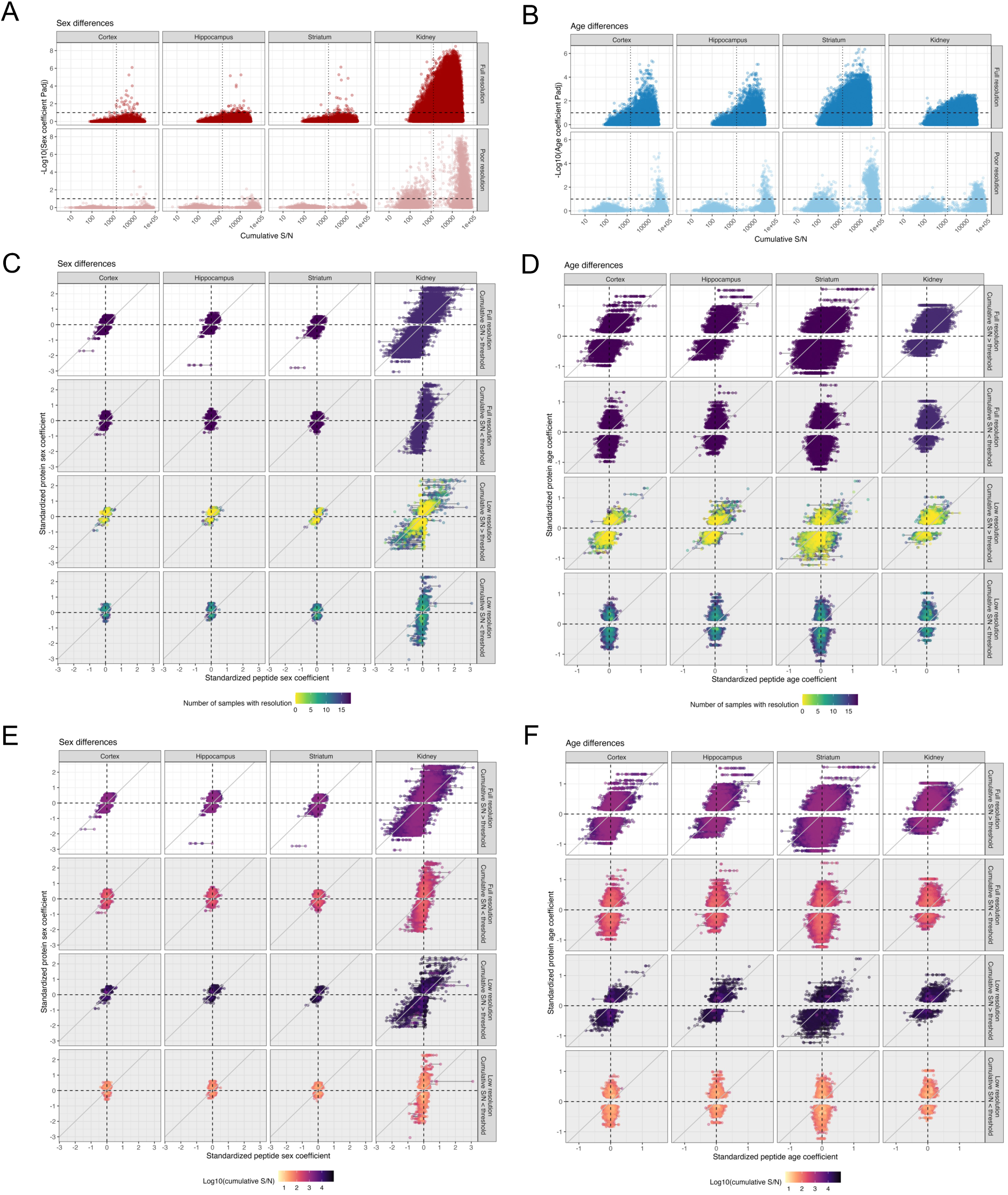
Impact of resolution and S/N on detecting age- and sex-differences in peptides. Association between statistical significance of test by cumulative S/N for (A) sex and (B) age differences, stratified by full resolution status. The vertical dotted line represents the cumulative S/N threshold for peptide filtering. Standardized regression coefficients for protein measurements by standardized regression coefficients from peptide measurements for (C, E) sex and (D, F) age, stratified by full resolution and cumulative S/N statuses. Peptides from the same protein are connected by lines. Points are colored by resolution (C, D) and cumulative S/N (E, F).

**Figure S2.**
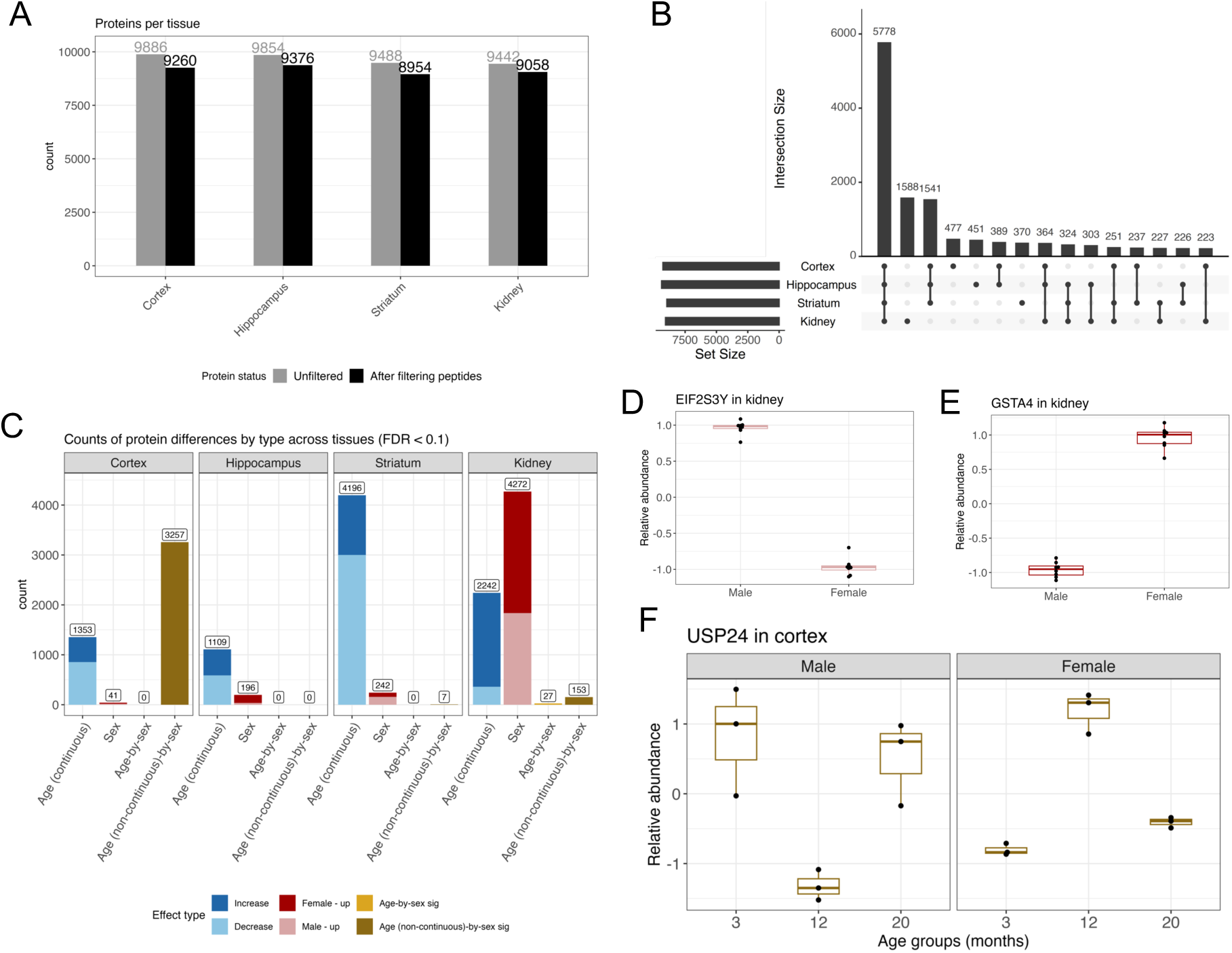
Summaries of protein counts and significant differences (FDR < 0.1). (A) Counts of detected proteins by tissues, before and after peptide filtration. (B) Upset plot summarizing the number of proteins observed across the four tissues. (C) Counts of statistically significant (FDR < 0.1) differences in each tissue. (D) EIF2S3Y had a sex difference in kidney, characterized by males having higher abundance. Data were scaled prior to plotting. (E) GSTA4 had a sex difference in kidney, characterized by females having higher abundance. Data were scaled prior to plotting. (F) USP24 had a non-continuous age-by-sex difference in cortex, characterized by flipped age-related abundance patterns between females and males. Data were scaled prior to plotting.

**Figure S3.**
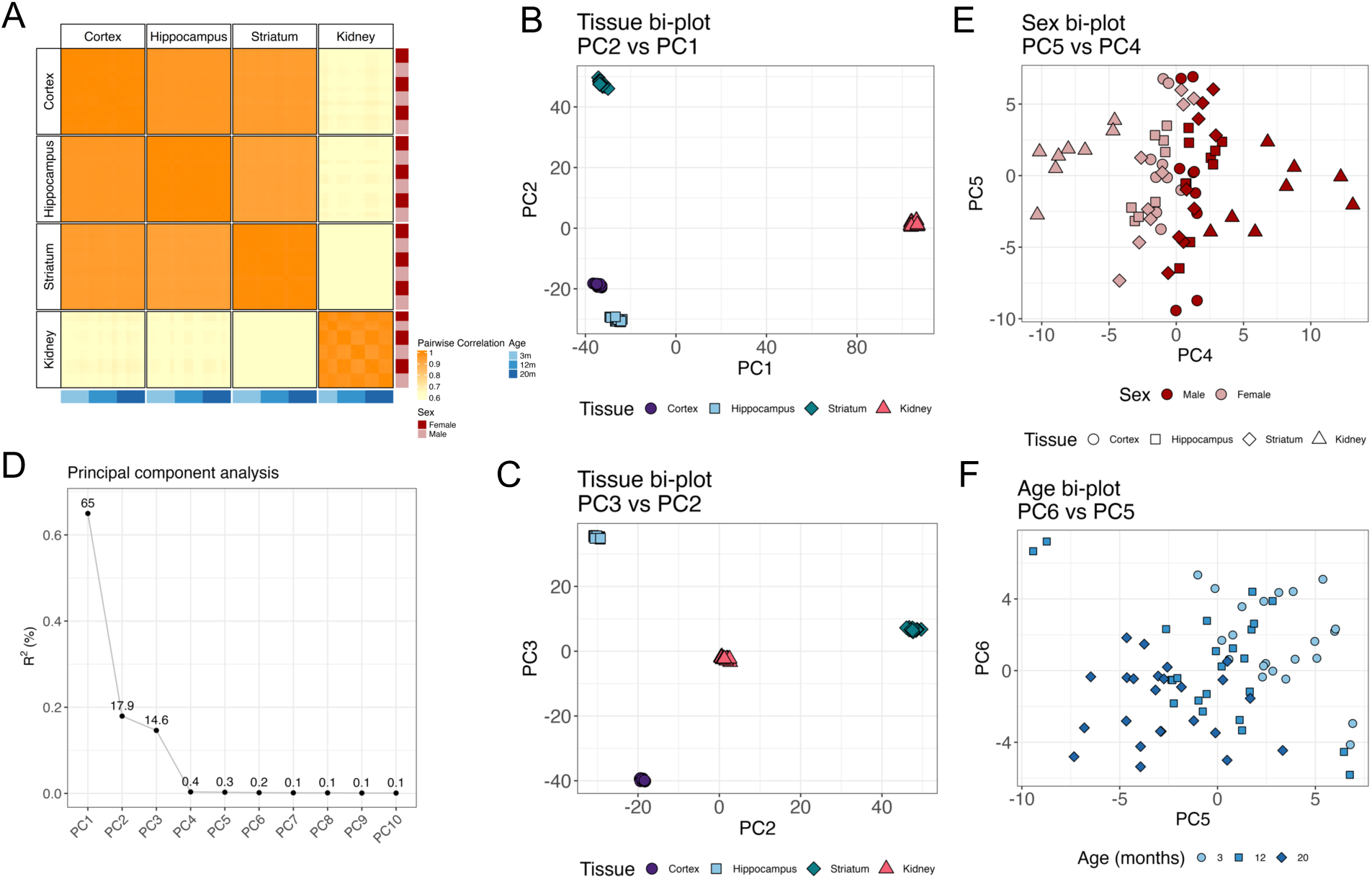
Principal component analysis reveals tissue, sex, and age as drivers of overall protein variation. (A) Heatmap of mouse-level correlation matrix based on the 5778 proteins observed across all four tissues. Columns and rows are ordered by sex and age groups to highlight potential contributions to correlation patterns. Bi-plots of (B) PC2 by PC1 and (C) PC3 by PC2, highlighting tissue as the major driver of variation across proteins. (D) Scree plot of the PCA, describing the proportion of overall variation explained by each PC. (E) Bi-plot of PC5 by PC4, highlighting that sex is the primary driver of PC4. (F) Bi-plot of PC6 by PC5, highlighting a continuous age effect across both PCs. PCA results for early development from Wang *et al.* 2024 is in Figure S7.

**Figure S4.**
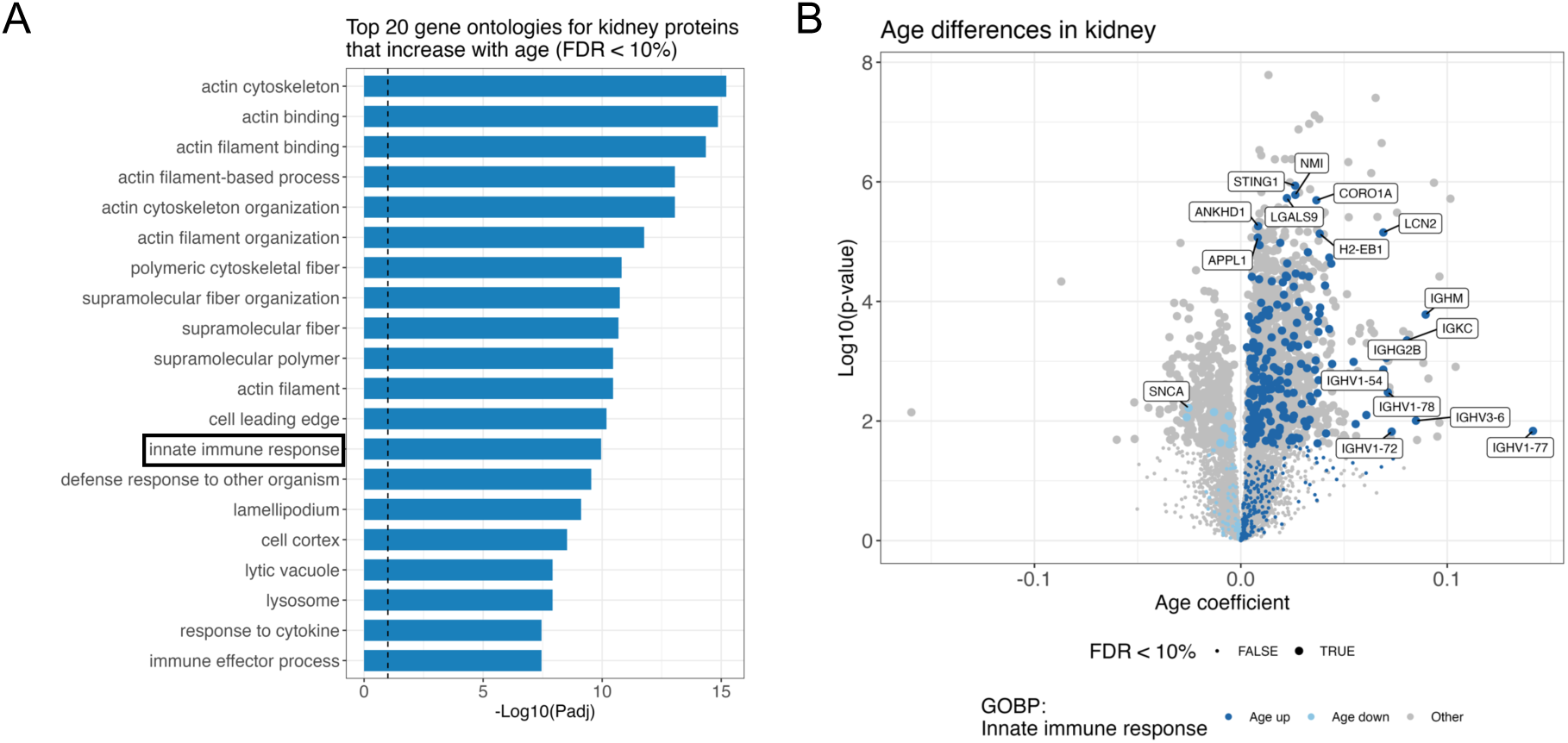
Gene set enrichments result for proteins that increase in abundance with age in kidney, including innate immune response proteins. (A) Top 20 gene ontologies enriched in kidney proteins with increased abundance with age (FDR < 0.1). Vertical dashed line represents a threshold of P_adj_ < 0.1. Innate immune response is highlighted with black box, which is featured in Fig. S4B. (B) Volcano plot of continuous age differences in kidney with innate immune response proteins highlighted.

**Figure S5.**
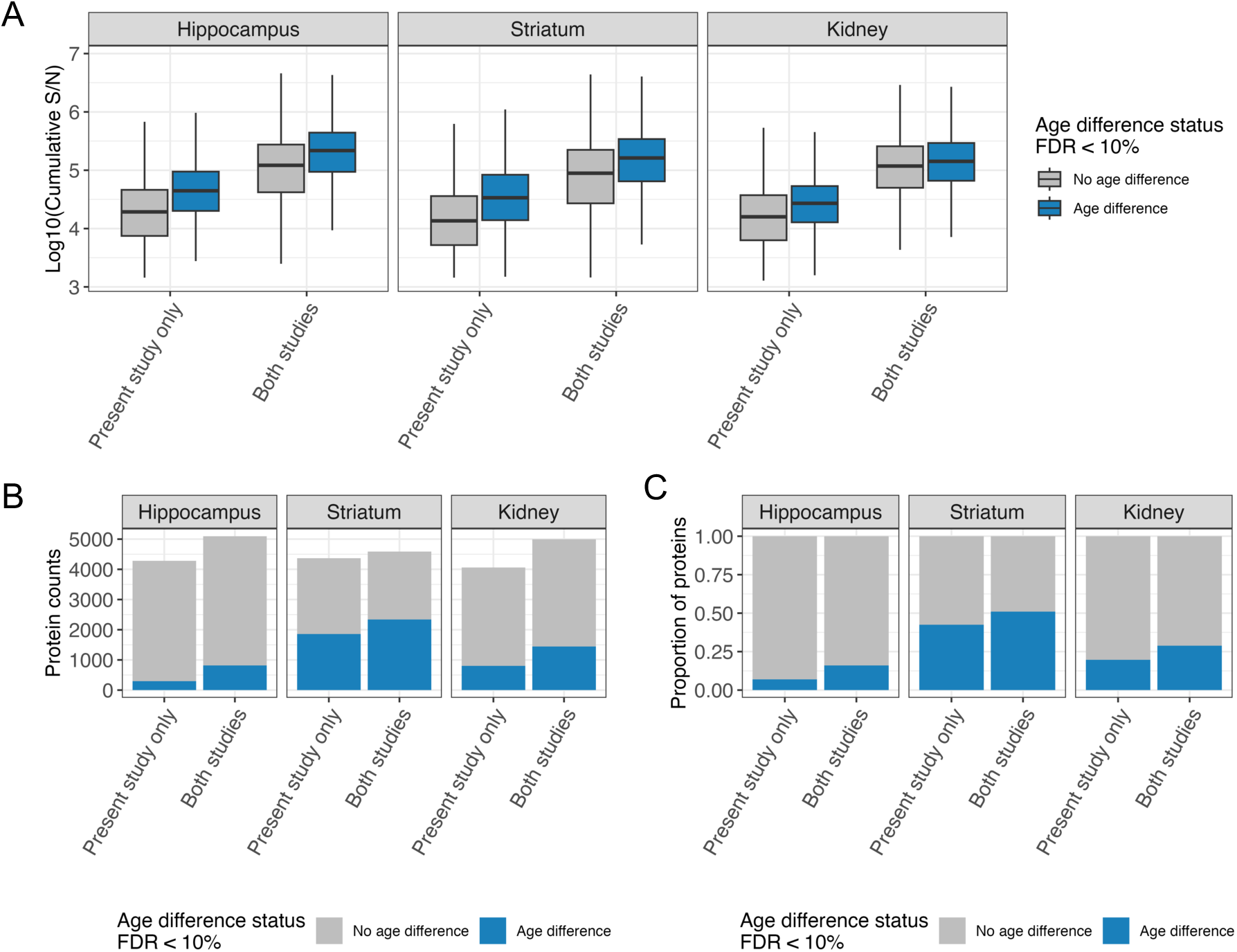
Breakdown of age difference detection for proteins based on whether they were observed in only the present study or in both present study and Keele *et al.* 2023. (A) Boxplots of cumulative S/N across proteins, categorized on whether they were observed in only the present study or both present study and Keele *et al.* 2023. (B) Count of proteins with and without significant age differences, categorized on whether they were observed in only the present study or both present study and Keele *et al.* 2023. (C) Proportion of proteins with and without significant age differences, categorized on whether they were observed in only the present study or both present study and Keele *et al.* 2023.

**Figure S6.**
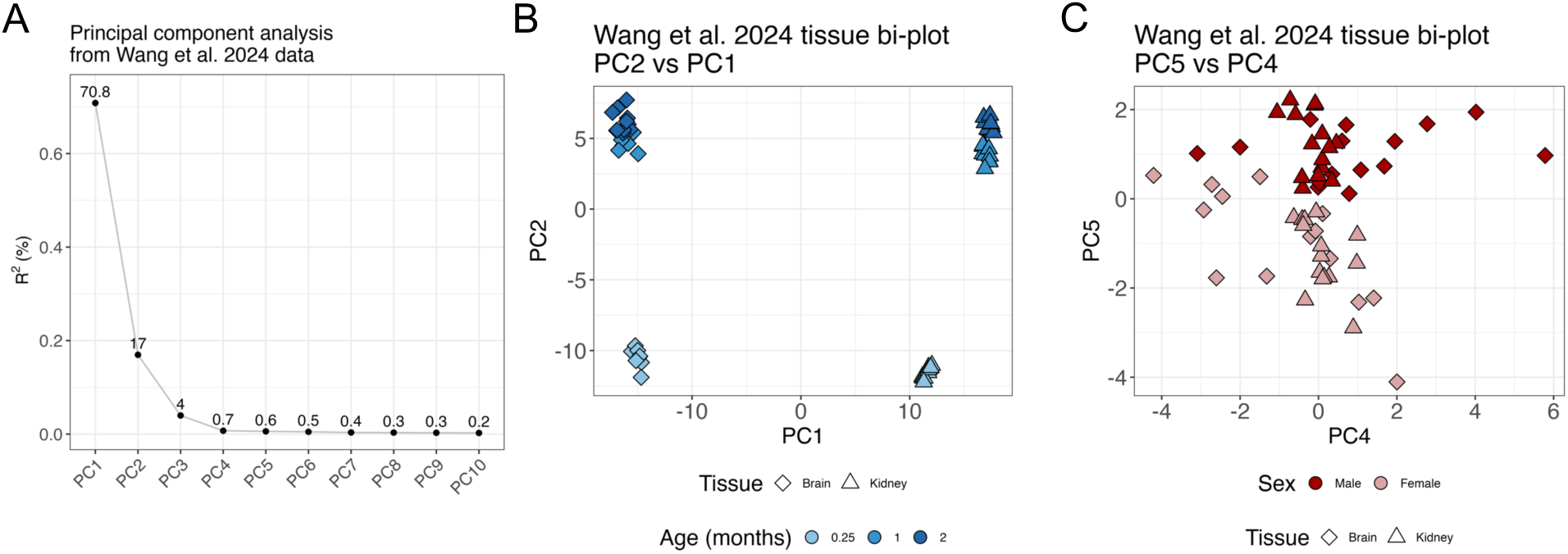
Principal component analysis reveals age in early development as a larger driver of overall protein variation than during adulthood. (A) Scree plot of the PCA, describing the proportion of overall variation explained by each PC. (B) Bi-plots of PC2 by PC1, highlighting tissue as the primary driver of PC1 and age as the primary driver of PC2. Variation due to age appears to be less continuous compared to adulthood aging (Fig. S3F), with the one-week age group being far separated from 1- and 2-month age groups. (C) Bi-plot of PC5 by PC4, highlighting sex as the primary driver of PC5. PCA results for adulthood aging from the present study are in Fig. S3.

**Figure S7.**
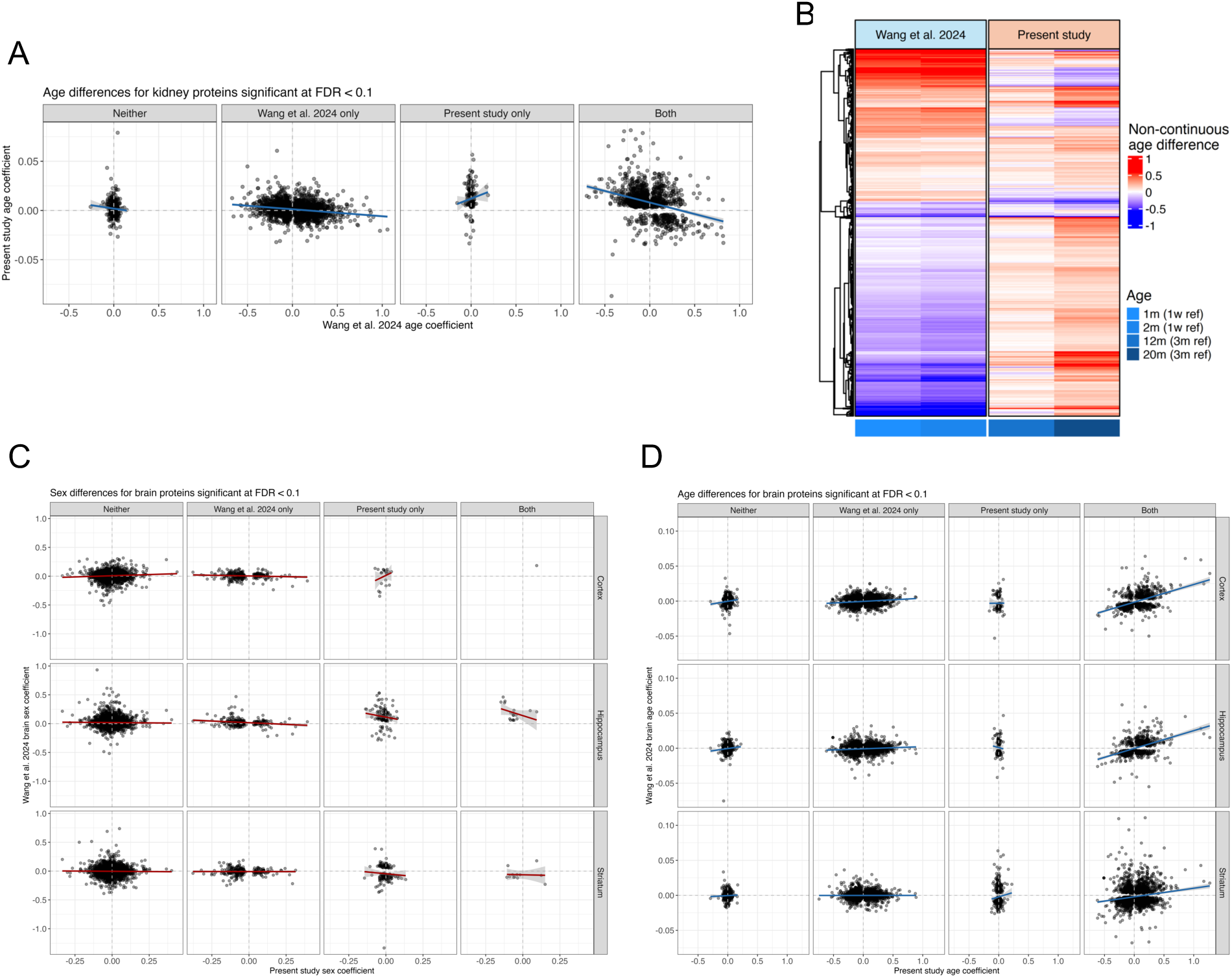
Comparison to early development protein data from Wang *et al.* 2024 reveals discordant aging patterns from adulthood depending on tissue. (A) Comparison of age difference coefficients for kidney between the present study and Wang *et al.* 2024. Best fit regression line and standard error interval included for reference. Coefficients are stratified into groups based on which studies detected a significant difference (FDR < 0.1). (B) Heatmap of non-continuous kidney age difference coefficients that were significant (FDR < 0.1) in both the present study and Wang et al. 2024. (C) Comparison of sex difference coefficients between three brain tissues from the present study and general brain tissue from Wang *et al.* 2024. Best fit regression line and standard error interval included for reference. Coefficients are stratified into groups based on which studies detected a significant difference (FDR < 0.1). (D) Comparison of continuous age difference coefficients between three brain tissues from the present study and general brain tissue from Wang *et al.* 2024. Best fit regression line and standard error interval included for reference. Coefficients are stratified into groups based on which studies detected a significant difference (FDR < 0.1).

